# Dietary iron deficiency impairs effector function of memory T cells following influenza infection

**DOI:** 10.1101/2024.07.22.604599

**Authors:** Marissa C. Bradley, Joshua Gray, Francesca La Carpia, Emma Idzikowski, Rebecca Guyer, Kalpana Pethe, Eldad A. Hod, Thomas J. Connors

## Abstract

The establishment of memory T cell responses is critical to protection against pathogens and is influenced by the conditions under which memory formation occurs. Iron is an essential micronutrient for multiple immunologic processes and nutritional deficiency is a common problem worldwide. Despite its prevalence, the impact of nutritional iron deficiency on the establishment of memory T cell responses is not fully understood. In this study we investigate the impact of nutritional iron deficiency on the generation, phenotype, and function of memory T cell responses using a murine model of dietary iron modulation in the context of influenza infection. Iron deficient mice have decreased systemic iron levels and develop significant anemia. Increased T cell expression of the transferrin receptor (CD71) is seen in iron deficient mice at baseline. During primary influenza infection, iron deficient mice experience increased weight loss and phenotypic evidence of impairments in T cell activation. Following recovery from infection, iron deficient mice generate increased influenza specific memory T cells which exhibit impaired ability to produce IFNγ, most notably within the lung. Importantly, the ability to produce IFNγ and TNFα is not recovered by co-culture with iron replete dendritic cells, suggesting a T cell intrinsic alteration in functional memory formation. Altogether, these results isolate a critical effect of nutritional iron deficiency on T cell memory development and function.

## INTRODUCTION

T cell mediated immunity is responsible for the eradication of dangerous pathogens and the development of long-lived memory responses. Iron is an essential micronutrient for many immunologic processes, including cellular activation and proliferation, which underlie the formation of protective T cell responses. Iron deficiency is a prevalent global problem frequently resulting from dietary insufficiency, particularly in children (1). Unfortunately, aberrant adaptive immune memory responses generated following primary encounters can result in decreased protection during subsequent exposures or in allergic responses that lead to chronic illnesses (2, 3). Individuals with iron deficiency are known to be more susceptible to infections and developing atopic diseases, suggesting a deleterious impact on immunity (4–6). Despite the worldwide prevalence of dietary iron deficiency, its effect on the generation and establishment of T cell mediated immunity has not been fully elucidated.

T cells have minimal labile iron pools and are largely dependent on uptake of iron from their environment (7–9). Genetic mutations in the transferrin receptor (CD71), responsible for mediating cellular uptake of iron, results in impaired cellular proliferation and reduced humoral immune responses, highlighting the critical role of iron in immune responses (10–12). Dietary iron deficiency leads to an overall reduction in T lymphocyte counts and thymic atrophy in mice (13). Interestingly, iron chelation has been shown to have differential impact on immune cell subsets (14, 15). Following iron chelation, T helper (Th) 1 cytokine production is reduced while Th2 functionality is largely unimpaired (14). Strikingly, hepcidin induced hypoferremia during primary infection or vaccination in mice results in reduced formation of antigen specific T cell responses and decreased ability to produce critical effector cytokines including IL-2, IFNγ and TNFα (16). Our current understanding of the role of iron in T cell mediated immunity is heavily reliant on genetic and chemical modulation models of iron availability. As nutrition is a leading cause of iron deficiency worldwide, isolating the role of dietary iron on the formation, maintenance, and function of memory responses following primary antigenic exposures is critical.

In this study we combine murine models of influenza infection and dietary iron modulation to reveal phenotypic and functional differences in memory T cell formation following infection. Differential systemic iron profiles and anemia develop in mice placed on iron deficient diets. Baseline immune profiling of splenic and lung tissue reveals similar numbers and phenotypes of T cell subsets with upregulation of CD71 noted in iron deficient mice. During primary infection iron deficient mice experience increased morbidity with increased and prolonged weight loss. Iron replete mice demonstrate evidence of earlier effector responses in the lung during acute infection which is associated with altered expression of activation makers CD25 and CD71. Following recovery, iron deficient mice had increased numbers of antigen specific T cells in the lung but decreased expression of markers required for the establishment of tissue residency. Strikingly, memory T cells formed during iron deficiency have decreased ability to produce inflammatory cytokines (IFNγ/TNFα) following stimulation. These results provide new insights into the role that iron plays in the generation of functional T cell immunity revealing tissue centric differences in memory T cell responses.

## MATERIALS AND METHODS

### Mice

Weanling female C57BL/6 mice were ordered from Charles River (Stone Ridge, NY) and housed in a pathogen-free BSL2 biocontainment facility. On day of arrival, mice were separated into two cohorts and provided with iron defined diets; replete (Envigo, Indianapolis IN; TD.110593, 45ppm Fe) or deficient (Envigo, TD.110592, 0-5ppm Fe) for the duration of the study. For all experiments, only mice on the same diet were cohoused (n = 5 per cage). For all studies both groups were euthanized at the same timepoints; 5 weeks on diet prior to infection, days post infection (DPI) 5 and 7 (5-6 weeks on diet), and DPI 28 (week 10 on diet). All animal studies were approved by the Columbia University Institutional Animal Care and Use Committee. All experiments were run in duplicate with representative data from a single replicate shown unless stated otherwise in figure legends.

### Hemoglobin Measurement

Starting at 4 weeks on iron defined diets, blood was drawn via tail-prick and collected into heparinized Eppendorf tubes. One mL of Drabkin’s reagent (Sigma-Aldrich, cat # D5941-6VL) was added to 4μl of blood, vortexed then kept in dark for 10 minutes. Samples were spun followed by absorbance reading at 540nm on Biotek ELISA reader (Agilent) along with standards prepared by serial dilutions of cyanomethemoglobin in Drabkin’s reagent (1:1 ratio). Standard curves generated were used to calculate hemoglobin concentration (g/dL).

### Iron-related measurements

All iron-related measurements were obtained at time of euthanasia. Non-heme iron was determined by a wet ashing procedure as previously described (17, 18). In brief, total liver weight was recorded then 100μg of liver tissue (wet weight) was dried overnight at 65°C. Liver tissue dry weight was recorded then 1ml of acid digestion mixture (3 M hydrochloric acid and 10% thiglycolic acid) was added and incubation continued for an additional 24 hours at 65°C. Samples were brought to room temp, centrifuged and 50μl aliquots were incubated with 200μl of chromogen (1.6M bathophenanthroline, 2M sodium acetate, 11.5M thiglycolic acid). Samples remained at room temperature for 30 minutes followed by measurement in duplicate of absorbance at 535nm using a microplate reader along with iron standards. Calculations of iron concentration for each sample were expressed as nanograms of iron per milligram of wet liver weight.

### Influenza infection

Mice were anesthetized by inhalation of isoflurane and infected intranasally with X31 (A/Hong Kong/1/68-x31 [H3N2]) influenza virus according to weight dose calculations. Each viral dose was consistent with 2000 TCID_50_ (median tissue culture infectious dose). Infection morbidity was determined by daily weight recordings and subsequent weight-loss at 30% or more was followed by immediate euthanasia.

### Tissue Processing

Following euthanizing, spleen tissue was mashed through a 100μm filter (Corning) and washed with phosphate buffered saline (PBS) (Fischer Scientific; MT21040CM). Lung tissue was dissociated using gentleMACS (Miltenyi Biotec) lung protocol and then digested for 1 hour in RPMI 1640 medium (Thermo Fisher; 11875093) containing collagenase D (Sigma-Aldrich), deoxyribonuclease (Sigma-Aldrich; COLLD-RO), and trypsin inhibitor (Sigma-Aldrich; 9002-07-7) in a 37°C shaking incubator. Samples were then filtered through a 100μm filter and washed in complete media (RPMI with 10% fetal bovine serum (FBS)). Red blood cell lysis was performed on all single cell suspensions using ACK lysis buffer (Thermo Fisher; A1049201) followed by wash in media.

### Flow Cytometry

Single-cell suspensions were resuspended in PBS with a 1:1000 dilution of Fixable Viability Dye for 30 min at 4°C. Cells were washed with fluorescence-activated cell sorting (FACS) buffer (phosphate buffered solution (PBS) with 5% heat-inactivated FBS) then resuspended in surface-stain antibody mix for 30 minutes at 4°C with a combination of the following antibodies; CD103-PerCPeFlour710 (Invitrogen clone 2E7), CD103-BUV563 (BD Biosciences clone M290), CD103-BV421 (Biolegend clone 2E7), CD11a-PerCPeFlour710 (Invitrogen clone M17/4), CD25-FITC (Biolegend clone PC61), CD25-PE (BD Biosciences clone PC61), CD3-AF700 (Biolegend clone 17A2), CD3-BUV805 (BD Biosciences clone 145-2C11), CD4-BUV395 (GK1.5), CD44-BV510 (Biolegend clone IM7), CD45-BV650 (Biolegend clone 30-F11), CD45-AF700 (Biolegend clone 30-F11), CD62L-BV605 (Biolegend clone MEL-14), CD69-PE/Dazzle594 or PE/Cy5 (Biolegend clone H1.2F3), CD71-PE/Cy7 (Biolegend clone R17217), CD71-BV750 (BD Biosciences clone C2), CD8a-BUV737 (BD Biosciences clone 53-6.7). Cells were washed with FACS-buffer and fixed with Fixation/Permeabilization Diluent (Invitrogen, ref 00-5223-56) and concentrate (Invitrogen, ref 00-5123-43) for 20 minutes at 4°C. Following fixation, cells were washed with FACS-buffer. For experiments including intracellular cytokine staining, samples were intracellularly stained in Permeabilization Buffer 10X (Invitrogen, ref 00-833-56) and deionized water for 30 minutes at 4°C with a combination of the following antibodies; IFNγ-BV750 (BD Biosciences clone XMG1.2), IFNγ-PerCP/Cyananine5.5 (Biolegend clone XMG1.2), IL17A-PE/Dazzle594 or FITC (Biolegend clone 18H10.1), TNFα-Pacific Blue (Biolegend clone MP6-XT22). Cells were washed and resuspended in FACS-buffer and flow cytometry data obtained on a Cytek (Fremont, California) Aurora 5L spectral flow cytometer or BD (Franklin Lakes, NJ) LSRII.

### Antigen-Specific Tetramer Staining

For X31 TCR-specific tetramer staining, single cell suspensions were incubated with CD4 IA_b_/NP_311-325_ and CD8 D_b_/NP_368-374_ tetramers (National Institutes of Health (NIH) Tetramer Core Facility (NTCF)) in 50μl at 37°C for 1 hour with intermediate agitation. Following incubation, additional surface markers were directly added, and the staining protocol follows the Flow Cytometry methods outlined above.

### T cell stimulations

Polyclonal stimulation: single cell suspensions were stimulated using PMA (Sigma, Cat#P1585) (50ng/mL) and ionomycin (Sigma, Cat#I9657) (1μg/mL) in a 96-well U-bottom plate in an incubator for 6 hours in the presence of GolgiStop (BD Biosciences, cat#554724) and GolgiPlug (BD Bioscience cat#555029). Following stimulation, cells were centrifuged, fixed then stained for intracellular cytokine detection (as described above).

Antigen specific stimulation: Influenza memory T cells were assessed for cytokine production as previously described (19, 20). Bone marrow was harvested from the hind legs of naïve iron replete mice. Bone marrow was centrifuged and resuspended in growth media (RPMI with GM-CSF (10ng/ml) and mouse Il-4 (20ng/mL)). Cells were added to a petri dish and transferred to the incubator (37°C, 5% CO_2_) for 7 days. Growth media was refreshed every 72 hours. Bone marrow-derived dendritic cells (BMDCs) were harvested at day 7 and re-plated at a concentration of 1×10^6^/ml in growth media. X31 (A/Hong Kong/1/68-x31 [H3N2]) influenza virus was sonicated then added to BMDC culture at a calculated multiplicity of infection of 0.3 and incubated overnight. The following morning unloaded (control) and antigen-loaded BMDCs were co-cultured with lung cells (100,000 BMDCs for every 5×10^6 lung cells) isolated from replete and deficient mice at DPI 28 post influenza infection. Net memory T cell cytokine production was determined by subtracting control from antigen-loaded co-culture assays.

### Statistical Analysis

Flow cytometry data were analyzed using FlowJo (ver. 10.10). Descriptive analyses and statistical testing were performed using Prism (GraphPad ver. 10.2.3).

### Materials availability

This study did not generate new unique reagents.

## RESULTS

### Combining nutritional and infection models to investigate T cell immunity

In order to investigate the role of isolated nutritional iron deficiency on the generation, phenotype, and functionality of memory T cells we leveraged our groups experience in mouse models of dietary iron restriction (21, 22) and influenza infection (23). In our combined model we placed weanling mice on iron defined diets (replete; 45ppm iron vs deficient; 0-5ppm iron, see methods) for the duration of the study. Decreased weight gain was observed in mice on iron deficient diets (Fig 1A). As significant iron restriction can induce anemia, we assessed hemoglobin levels as a surrogate marker of iron status starting at the 4^th^ week on defined diets. Mice on iron deficient diet developed anemia with significantly decreased hemoglobin compared to replete mice (Fig 1B). To confirm iron status prior to immunologic assessment, we quantified total iron levels in the liver (see methods). Mice on the deficient diet had significantly reduced iron levels (Fig. 1C), similar to our previous experience (21, 22). The confirmation of distinct iron profiles between the two groups provided the ability to assess the impact of iron deficiency on T cell immunity.

**Figure 1:**
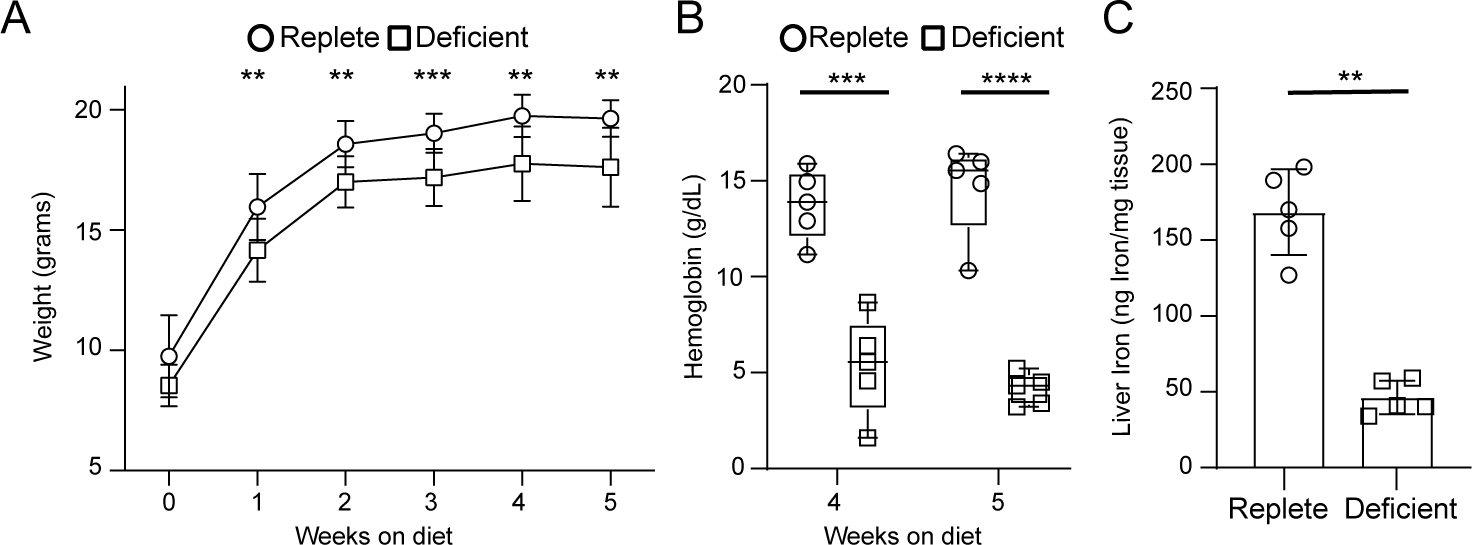
Mouse model of dietary iron deficiency. A) Weight curves by weeks on iron defined diets for each cohort, boxes represent mean with bars representing standard deviation. B) Comparison of hemoglobin by group at weeks 4 and 5 on defined diets. Bars represent mean and range with boxes depicting interquartile range. C) Quantification of liver iron at diet week 5 for each cohort, box with bars represents mean with standard deviation. For all panels, n=5 mice per group. All statistical testing via Mann-Whitney t-test. ** = p<0.01, *** = p <0.001, **** = p <0.0001.

### Iron deficient T cells express increased levels of the transferrin receptor at baseline

We sought to determine if nutritional iron deficiency impacted T cell parameters at baseline in naïve mice using flow cytometry (Fig. S1A). Replete and deficient mice had similar total number of T cells in the spleen, but decreased T cells were seen in lung tissue of deficient mice (Fig. 2A). Subtyping of T cells into naïve (CD44^−^/CD62L^+^) and CD44^+^ subsets revealed decreased proportions of naïve T cells in the spleen of deficient mice, which were not detected in the lung (Fig. 2B). We next examined systemic (splenic) T cells for differences in phenotypic expression of markers associated with activation (CD69) or antigenic experience (CD11a). We found no difference in proportions of CD69 and CD11a expression within non-naïve T cell subsets between the two groups (Fig. 2C). As iron is transported into the cell by the transferrin receptor (CD71), we analyzed T cell expression of CD71 between the groups. Increased expression of CD71 was seen in bulk T cell populations from iron deficient mice with significant differences driven by CD71 expression on non-naive CD4^+^ and CD8^+^ T cells (Fig. 2D). These results reveal differences in T cell phenotypes in the setting of nutritional iron deficiency, which could lead to varying T cell responses upon antigenic challenge.

**Figure 2:**
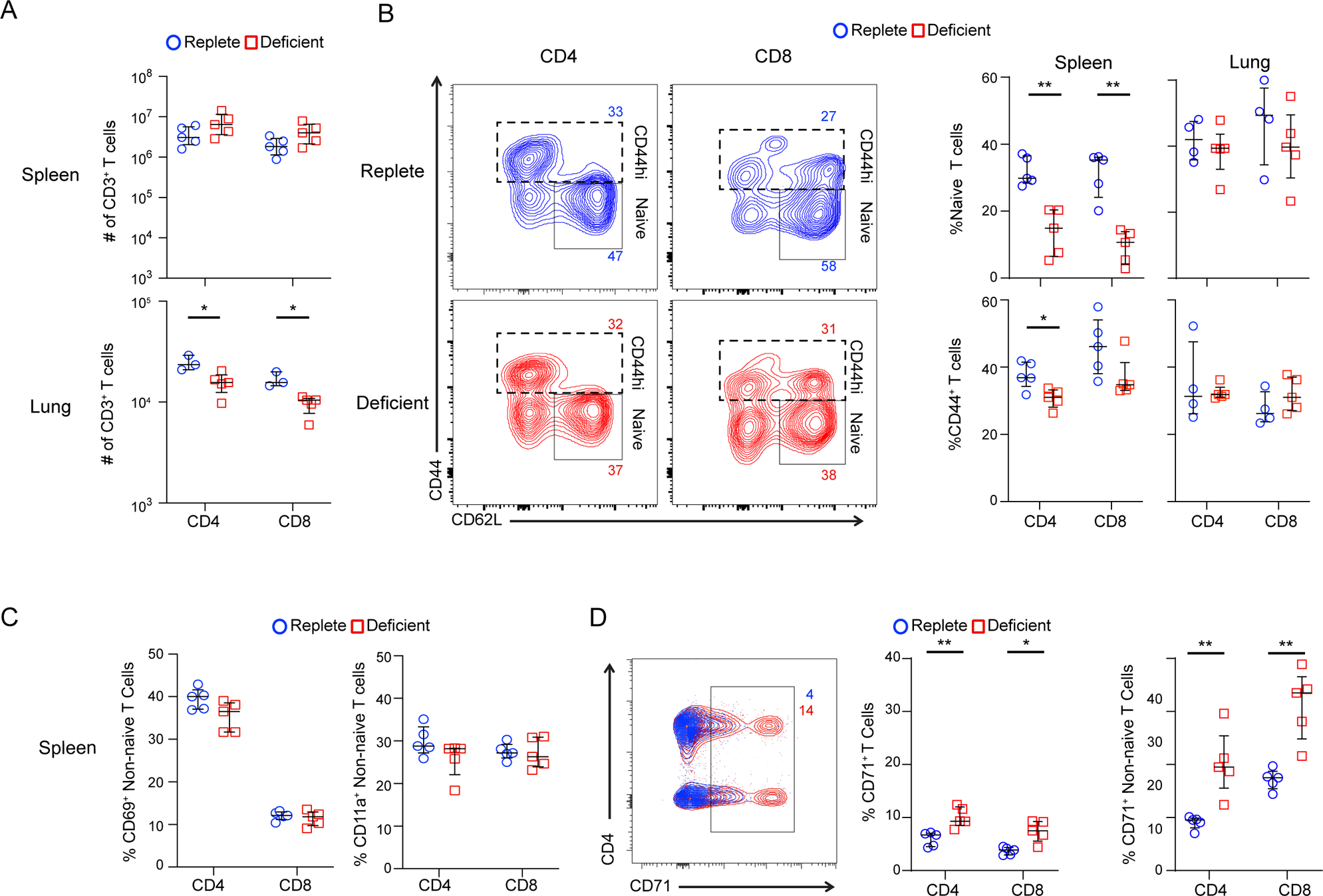
Phenotypic differences in T cell profiles at baseline across iron cohorts. Flow cytometry analysis of naïve mice T cells at week 5 on iron defined diets. A) Counts of CD4^+^ and CD8^+^ T cells in spleen (top) and lung (bottom) by cohort. Lines represent mean with standard deviation. B) Analysis of memory markers on CD4^+^ and CD8^+^ T cells (naïve; CD44^−^/CD62L^+^) within representative lung tissue (left) and compiled data (right) organized by tissue site. Lines represent mean with standard deviation. C) Analysis of surface marker expression (CD69; left and CD11a; right) on splenic non-naïve T cells. Lines represent mean with standard deviation. D) Analysis of transferrin receptor (CD71) expression on splenic T cells with representative data (left), and compiled data for expression on all T cells (middle) and non-naive T cells (right). Lines represent mean with standard deviation. For all panels, n= 3-5 mice per group. Statistical testing by Mann-Whitney t-test. * = p<0.05, **=p<0.01.

### Increased morbidity and differential phenotypes associated with iron deficiency during primary infection

The phenotypic differences observed at baseline led us to investigate the role of iron following viral respiratory tract infection. Mice were intranasally infected with influenza (X31, see methods). Mice with iron deficiency were noted to have increased weight loss at early time points (DPI 4 and 5) compared to replete mice (Fig. 3A). Iron replete mice began to regain weight earlier (DPI 6/7), coinciding with the beginning of a typical peak T cell response (23, 24), whereas iron deficient mice continued to lose weight until DPI 8 and exhibited slower recovery of weight (Fig. 3A). We examined local and systemic T cell responses at DPI 5 and 7 to determine if differences in T cell responses existed. Total T cell counts were similar in the lung for both groups at DPI 5 and 7 but increased numbers of both CD4^+^ and CD8^+^ T cells were identified in the spleen at DPI 5 of iron deficient mice (Fig 3B). We examined the expression of CD44 to explore differences in early effector and memory subset responses. Decreased proportions of CD8^+^ CD44^+^ cells were seen in the lung of deficient mice earlier in infection (Fig. 3C). We next explored phenotypic markers (CD25 and CD71) known to be directly correlated with activation and proliferation of T cells (25). We found differential expression of CD25 on T cells in spleen and lung (Fig. 3D). Iron replete mice largely showed their highest expression at the earlier time point (DPI 5), whereas deficient mice showed increasing or stable levels of expression between DPI 5 and 7 (Fig. 3D). Iron deficient mice demonstrated persistently increased expression of CD71 at later timepoints contrasted to the falling expression between DPI 5 and 7 exhibited by replete mice (Fig. 3E). As the generation of antigen specific responses are critical to pathogen clearance and memory establishment, we used tetramer staining (Fig. S1B, see methods) to identify influenza specific T cells. Consistent with the differences seen in activation, we found higher numbers of CD8^+^ influenza specific cells in the lung of iron replete mice at DPI 7 (Fig. 3F). Combined, the alterations in proportions of effector cells and activation markers suggest a deleterious impact of iron deficiency on localized primary T cell responses.

**Figure 3:**
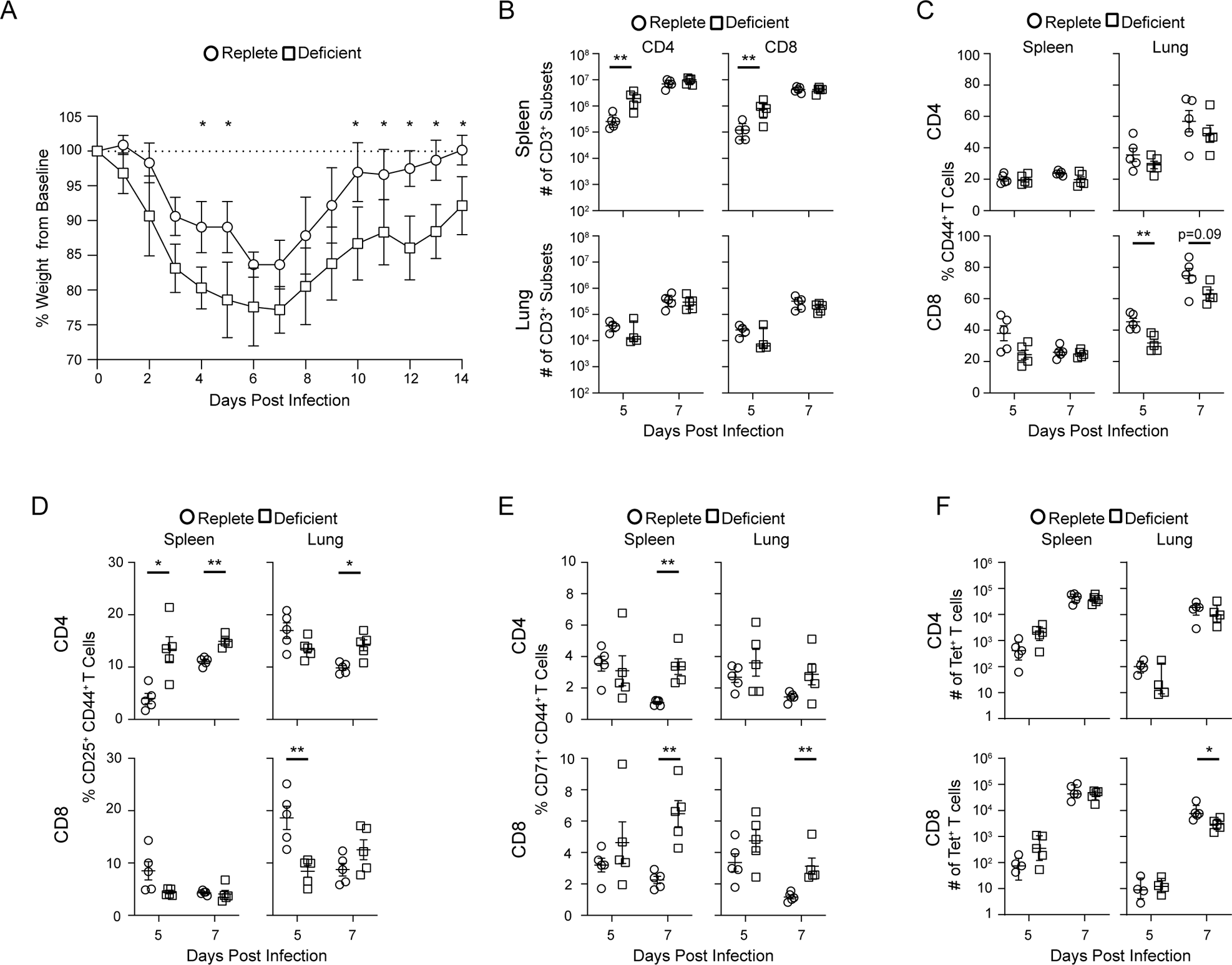
Increased morbidity with differential T cell responses in iron deficient mice following primary influenza infection. Assessment of differences related to iron status on T cell responses during acute influenza infection. Mice infected at week 5 on iron defined diets. A) Weight (%) from baseline by day post infection (DPI) following influenza (X31) infection. Data shown as mean with standard deviation. B-E) Flow cytometry analysis of T cell subsets during acute infection, at DPI 5 and 7 organized by tissue and subset. Bars represent mean with standard deviation. B) Counts of CD3^+^ T cells, C) Analysis of CD44 expression on T cells, D) CD25 expression on CD44^+^ T cells, E) CD71 expression on CD44^+^ T cells. F) Counts of antigen specific T cells at DPI 5 and 7 organized by tissue location and T cell subset as determined by tetramer staining. Bars represent mean with standard deviation. All statistical testing performed with Mann-Whitney t-test. All experiments with data derived from 3-5 mice per group. * = p<0.05, ** = p<0.01.

### Iron deficiency results in decreased anti-viral functionality in CD8**^+^** T cells

The differences in T cell phenotypes during acute infection led us to investigate whether the generation and function of memory T cells was impacted by iron deficiency. We analyzed T cell responses in mice which had recovered from influenza infection at DPI 28. Total CD4^+^ and CD8^+^ T cell numbers were similar in both lung and spleen across groups (Fig. 4A) with increased proportions of CD4^+^ effector memory (T_EM_; CD44^+^/CD62L^−^) phenotype seen in lung tissue of iron deficient mice (Fig. 4B). A subset of T_EM_ are known to establish residency within the tissue environment, designated as tissue resident memory T cells (T_RM_), and provide optimal protection upon re-exposure (26–28). We examined phenotypic markers associated with T_RM_ (CD69 and CD103) and found decreased proportions of CD8^+^ T_RM_ within the lung of iron deficient mice, suggesting a skewed memory T cell compartment (Fig. 4C). Interestingly, we identified increased accumulation of influenza specific CD4^+^ and CD8^+^ T cells in the lung of iron deficient mice (Fig. 4D). We performed intracellular cytokine staining following PMA/Ionomycin stimulation, which reveals T cell intrinsic function by bypassing the T cell receptor (Fig 4E and see methods). Within the spleen, T cells showed similar ability to produce key effector cytokines (IFNγ and IL-17a) (Fig. 4E). Strikingly, within the lung tissue both CD4^+^ and CD8^+^ T cells of deficient mice demonstrated significantly decreased capacity to produce IFNγ (Fig. 4E), suggestive of a T cell intrinsic defect.

**Figure 4:**
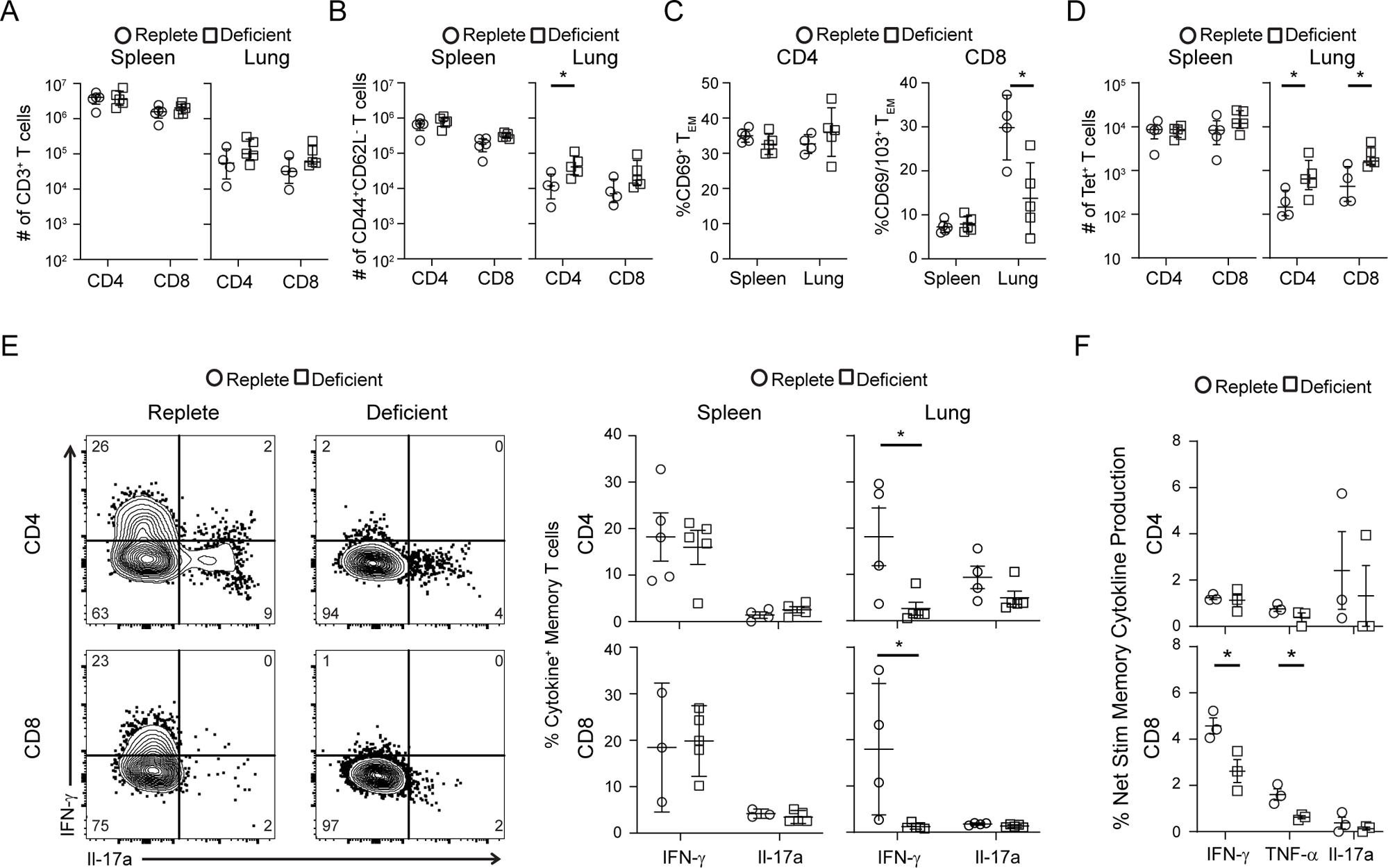
Altered phenotype and function of lung memory T cells generated under iron deficient conditions. Flow cytometric analysis of memory T cell phenotypes and intracellular cytokine production at 28 days post influenza infection (9 weeks on iron defined diets). A) Counts of T cell subsets from spleen and lung. Bars represent mean with standard deviation. B) Proportions of T effector memory (T_EM_; CD44^+^/CD62L^−^) cells in spleen and lung. Bars represent mean with standard deviation. C) Analysis of residency markers CD69 (CD4^+^) and CD69/CD103 (CD8^+^) on T_EM_ populations for spleen and lung. Bars represent mean with standard deviation. D) Assessment of influenza specific T cells by tetramer staining. Bars represent mean with standard deviation. A-D) n = 4-5 mice per group. E) Left; Representative data for intracellular cytokine staining from CD44^+^ T cell subsets following PMA/Ionomycin stimulation, organized by diet cohort. Right; Compiled data organized by tissue and subset. F) Compiled data from intracellular cytokine staining of T cells following co-culture with influenza antigen loaded iron replete antigen presenting cells (n=3 mice per group) with results of single experiment shown. For all panels, n= 3-5 mice per group. All statistical testing with Mann-Whitney t-test. * = p<0.05.

Across multiple experiments, the lung tissue was the site where T cell responses were most different between the two groups. We speculated that the spleen may offer more available iron (29) than the lung and that functional differences previously observed (Fig. 4E) would be reversible. To remove the impact of local iron availability and antigen presentation, we harvested bone marrow from replete mice to differentiate into iron-replete BMDCs. BMDCs were loaded with influenza and co-cultured with T cells isolated from the lungs of replete and deficient mice recovered from influenza infection (DPI 28). We then assessed their ability to produce cytokines (Fig. 4F and see methods). Intracellular cytokine staining again revealed decreased production of key anti-viral cytokines IFNγ and TNFα by CD8^+^ T cells from iron deficient mice (Fig. 4F). Altogether, these data reveal that T cell memory generation occurs in the setting of dietary iron deficiency, but phenotype and function is significantly affected.

## DISCUSSION

This study identifies derangements in anti-viral effector and memory T cell responses in the setting of isolated dietary iron deficiency. Nutritional iron deficiency leads to decreased systemic iron stores, anemia, and increased expression of CD71 on T cells in iron deficient mice. During primary influenza infection, iron deficient mice experienced increased morbidity and evidence of impaired T cell activation, particularly in the local pulmonary environment. In recovery, iron deficient mice have increased numbers of influenza specific T cells but altered phenotypes. Strikingly, memory T cells have decreased ability to produce key anti-viral cytokines and this functionality is not restored when cultured with iron replete DCs. These findings are most apparent within the lung tissue and suggest a defect in localized memory T cell formation and functionality as a result of iron deficiency. This study provides new insights into the impact of dietary iron deficiency on primary and memory T cell responses.

Primary immune responses are critical events for the eradication of pathogens and establishment of future protective immunity. The metabolic demands of T cells which are undergoing activation and differentiation are vastly different from those at rest, requiring shifts in programming and nutrient usage (30, 31). Deprivation of key nutrients results in altered T cell activation and effector function (32–34). Iron is a critical co-factor in many immunologic processes and T cells are particularly susceptible to deficiency (13, 16, 35). In our study, iron deficient T cells demonstrated delayed activation during acute infection which was associated with increased morbidity and delayed recovery. These effects were most notable within the lung environment. Localized immune responses, which are critical for optimal protection from pathogens, are influenced by their environment and can lead to allergic responses based on the conditions under which they are formed (26, 36, 37). In children with malnourishment, T cells display reduced expression of activation markers (CD69/CD25) and a shift in effector function with reduced T helper 1 (IFNγ and IL-2) cytokines and increased allergic Th2 (IL-4) cytokine production (34). Whether iron deficiency skews T cell responses towards more allergic responses at the expense of inflammatory function warrants further exploration. Additionally, as both protective and allergic response are highly tissue centric, improved understanding of how iron impacts T cell immunity, particularly within the tissue environment, is needed to delineate the role of iron in immunologic health.

Mature memory responses develop phenotypic and functional adaptations to their tissue environment (38). The most dynamic period of memory T cell generation and development occurs over the course of infancy and childhood (39). Infancy and childhood also correlate to periods of increased iron requirements with nutritional and dietary restrictions being a leading cause of iron deficiency in this age group (40). We show that dietary iron deficiency results in impaired inflammatory effector function of memory T cells which was not reversed by providing optimal recall conditions. This suggests that memory formed during periods of iron deficiency could lead to poor protection upon repeat antigenic exposures and may provide an immunologic link to the association of iron deficiency with allergic and atopic responses (3, 6, 41). We were not able to fully assess the impact of iron on antigen presentation or the tissue environment, which play essential roles in adaptive immunologic memory (42). Further studies dissecting the mechanisms of immune cell interactions and the long-term impacts of memory responses formed during periods of iron deficiency would help to elucidate avenues for intervention. As infancy and childhood are dynamic periods of immune development with significant implications for decades of health, interventions during this period could have outsized effects.

Globally, iron deficiency is a prevalent problem associated with increased morbidity during primary antigenic exposures and reduced responsiveness to vaccinations (43–45). Improving iron status at time of vaccination has shown promise in improving immune outcomes in children (43). As correction of iron status is a potentially addressable health issue, additional research that focuses on the role of iron in adaptive immunologic memory formation, identifies critical levels which warrant intervention, and defines windows of opportunity for intervention are needed. In summary, our results provide new insights into the fundamental role of iron in T cell immune responses, which are essential for tissue health and durable protection against pathogens.

## ACKNOWLEDGEMENTS

We thank Donna Farber, PhD and Gary Brittenham, MD for their guidance and insight. We thank Isaac Jensen, PhD for critical reading of this manuscript.

## FOOTNOTES

### 2. Author Contributions

Conceptualization, M.C.B., E.H. and T.J.C.; data curation, M.C.B., and T.J.C.; formal analysis, M.C.B., and T.J.C.; funding acquisition, T.J.C.; investigation, M.C.B., J.G., F.L., E.I., R.G., K.P., E.H., and T.J.C.; methodology, M.C.B., J.G., F.L., E.H., and T.J.C.; project administration, T.J.C.; supervision, T.J.C.; visualization, M.C.B., and T.J.C.; writing - original draft, M.C.B. and T.J.C; writing - review & editing, all authors.

