## Supplemental Figure 1 for "Dietary iron deficiency impairs effector function of memory T cells following influenza infection"

Figure S1

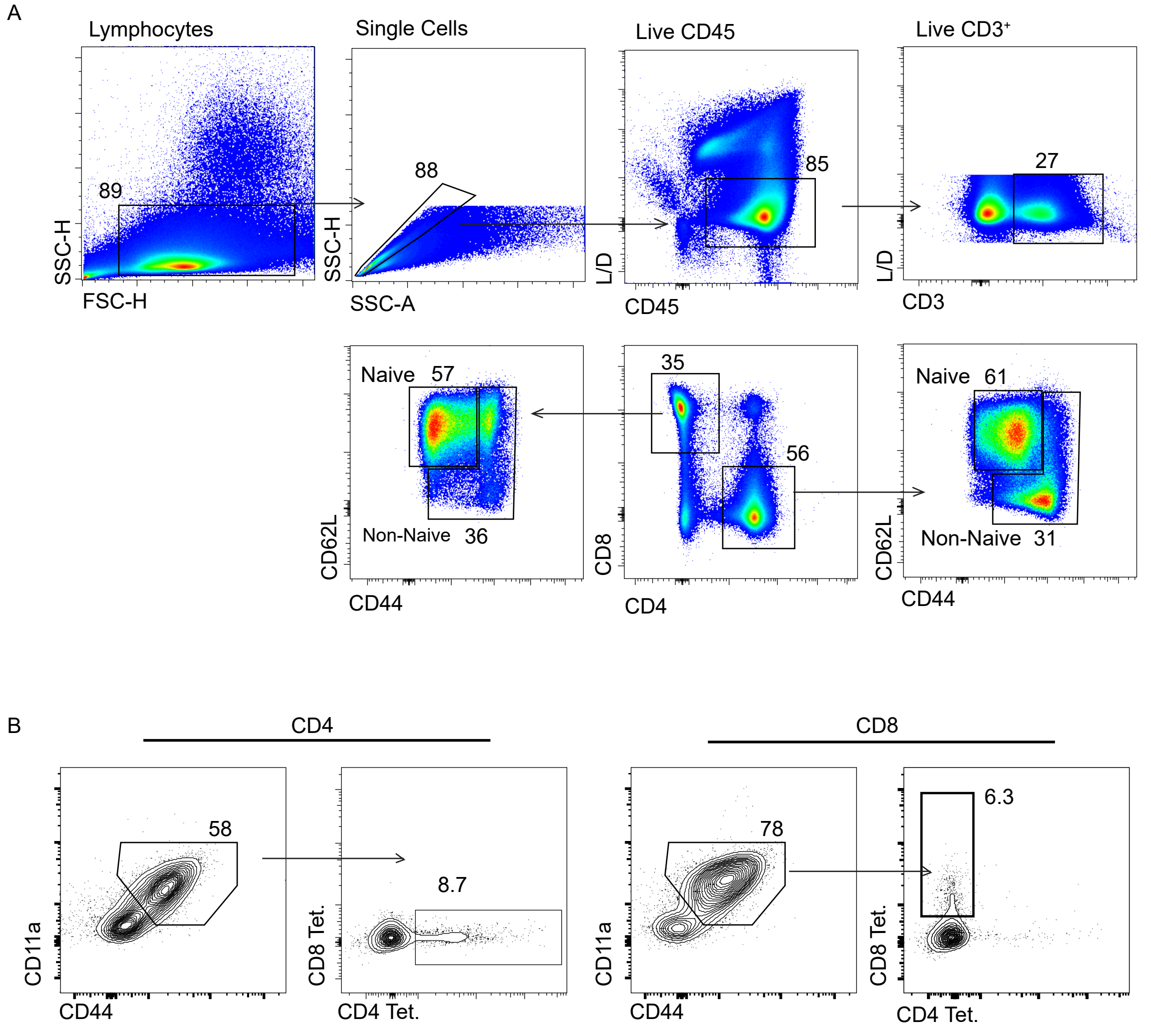


Figure S1: Flow cytometry gating strategy. A) Live, CD45⁺, CD3⁺ cells were gated by CD4 and CD8 expression then CD44 and CD62L to define memory subsets. B) Antigen experienced T cells (CD11a⁺/CD44⁺) gated by expression of influenza tetramers (Tet.).
